# Single-Cell Transcriptomics Reveals Tumorigenic Lineages and Interactions Persisting and Evolving From Premalignancy to Cancer in the Stomach

**DOI:** 10.64898/2026.11.04.764362

**Authors:** Emily N. Vazquez, Clarisel C. Lozano, Xiaodian Sun, Dyana Torres Juvier, Chris M. Li, Wael El-Rifai, Alexander Zaika, Yuguang Ban, Yan Guo, Xiaodong Cai, Zhibin Chen

**Affiliations:** Department of Microbiology and Immunology, University of Miami Miller School of Medicine, Miami, FL, USA; Sylvester Comprehensive Cancer Center, University of Miami, Miami, FL, USA; Division of Biostatistics and Bioinformatics, Department of Public Health Sciences, University of Miami Miller School of Medicine, Miami, FL, USA; Department of Surgery, University of Miami Miller School of Medicine, Miami, FL, USA; Department of Veterans Affairs, Miami Healthcare System, Miami, FL, USA; Department of Electrical and Computer Engineering, University of Miami, Miami, FL, USA

**Keywords:** Embigin, Fibronectin, metaplasia, chief cells, gastric cancer

## Abstract

**Background & Aims:** Gastric cancer is often diagnosed at advanced stages, making the premalignant window critical for early detection. However, the epithelial changes that characterize this stage remain poorly defined. We used CTLA4-knockdown (CTLA4KD) mice, which model human CTLA4 insufficiency-associated gastric cancer, that develops autoimmune gastritis, metaplasia, and adenocarcinoma, to define molecular changes across disease progression.

**Methods:** Single-cell RNA sequencing and 40-color spectral flow cytometry were performed on gastric tissue across premalignant and cancer stages. We evaluated epithelial lineage relationships, cell-cell communication pathways, and candidate markers at the transcript and protein levels in murine and human gastric adenocarcinoma.

**Results:** We identified a metaplastic/tumorigenic population in CTLA4KD mice that emerged in premalignancy with a chief cell transcriptomic origin. Cell-cell communication analysis revealed a progressive increase in fibronectin (FN1) signaling from premalignancy through cancer, transitioning from stromal to autocrine signaling in the metaplastic/tumorigenic population. Expression of Embigin (EMB), a fibronectin receptor, increased at both transcript and protein levels in premalignancy and remained elevated in cancer. EMB⁺ epithelial cells co-expressed CD133, a cancer stem cell marker, consistent with a stem-like phenotype. EMB expression was also higher in human gastric adenocarcinoma than in matched normal tissue.

**Conclusions:** Using a model that captures the full histopathological cascade of gastric tumorigenesis, we delineate single-cell transcriptional programs that persist and evolve from premalignancy to adenocarcinoma. In-depth computational analyses identified novel markers involved in cell-cell communication during tumorigenesis, including EMB, an epithelial marker that persists from premalignancy through cancer in the murine model and validated in human gastric adenocarcinoma, suggesting EMB as a marker of premalignant progression.

**Graphic Abstract:** 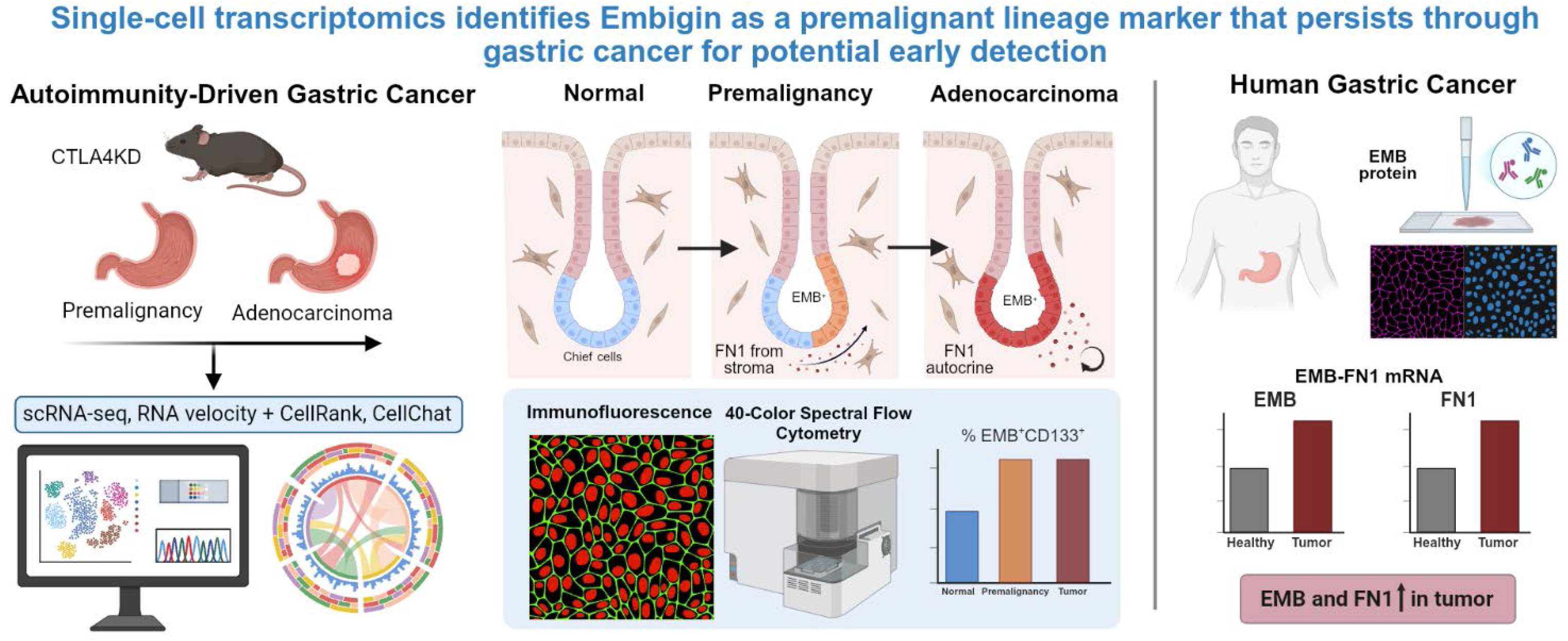

**What You Need to Know:** *Background and Context:* The cellular and molecular changes that drive gastric premalignancy toward invasive cancer remain poorly defined, leaving no reliable way to identify which premalignant lesions will progress.

*New Findings:* We identified a tumorigenic epithelial population present in premalignancy and persisting into cancer, accompanied by increased fibronectin signaling and sustained Embigin expression across disease progression.

*Limitations:* We defined the premalignant-to-cancer transition in a mouse model and validated a marker in human gastric adenocarcinoma. Single-cell studies of human gastric premalignancy to cancer progression are needed to confirm these findings.

*Clinical Research Relevance:* Defining epithelial changes that persist from premalignancy through cancer may help distinguish lesions progressing toward malignancy from those remaining stable. Embigin may represent a candidate marker of this transition, supporting future studies aimed at improving risk stratification and earlier identification of patients most likely to develop gastric cancer.

*Basic Research Relevance:* This study defines cellular and molecular changes during the transition from gastric premalignancy to cancer. It identifies a tumorigenic population arising from chief cells that persists across disease stages, as well as changes in fibronectin signaling and Embigin expression associated with progression of premalignancy toward invasive cancer.

*Lay Summary:* We identified markers of stomach cells on the pathway to becoming cancer cells in animal models, and these markers may provide a tool for early detection before invasive cancer develops.

## Introduction

Gastric cancer is the fifth most common malignancy worldwide^1^. Adenocarcinoma accounts for more than 90% of gastric cancer cases and is thought to develop through a sequence of histopathological changes classically described by the Correa cascade, in which chronic inflammation drives the progression from normal mucosa through atrophy, intestinal metaplasia (IM), and dysplasia to invasive adenocarcinoma^2^. A more recently identified histopathological change along this cascade, preceding or coexisting with IM, is spasmolytic polypeptide-expressing metaplasia (SPEM), a mucous-cell metaplasia that arises from the reprogramming of the gastric epithelium and is increasingly recognized as a precursor to gastric cancer^3, 4^. Although the inflammatory initiator has been mainly attributed to *Helicobacter pylori* infection^5^, the epidemiology of gastric cancer has shifted in recent years. *H pylori* infection rates are declining in developed nations, yet gastric cancer diagnosis among young individuals continues to rise^6^.

This trend parallels the rise in other early-onset malignancies linked to inflammatory and autoimmune conditions^7–9^. Among these, autoimmune gastritis is recognized as an etiologic factor for gastric cancer^10, 11^. In autoimmune gastritis, autoreactive T cells target the parietal cell H^+^/K^+^ ATPase, causing parietal cell loss, gastric atrophy, and a predisposition to tumorigenesis^12^. Consistent with an autoimmune contribution, gastric cancer diagnoses are rising faster in young women than in young men^13^, possibly related to the disproportionate prevalence of autoimmune diseases in females^14, 15^.

Autoimmunity arises from a breakdown of immune tolerance, a process in which the immune checkpoint CTLA4 plays a major role. Yet CTLA4 presents a paradox in gastric cancer. Although CTLA4 is a canonical inhibitor of antitumor immunity, reduced CTLA4 activity increases gastric cancer risk; patients with CTLA4 haploinsufficiency develop atrophic gastritis and gastric cancer^16, 17^, and gastric cancer-associated CTLA4 polymorphisms reduce CTLA4 expression^18–21^. These findings implicate CTLA4 insufficiency in gastric cancer risk, likely through chronic autoimmune inflammation arising from impaired immune tolerance.

Our group previously generated a transgenic CTLA4 shRNA knockdown mouse model (CTLA4KD) in which CTLA4 expression is reduced by approximately 60%, recapitulating the CTLA4 reduction seen in patients with haploinsufficiency. By 5-6 weeks of age, these mice develop autoimmune gastritis and SPEM, and after 12 months they progress spontaneously to invasive gastric cancer^22^. To our knowledge, this is the only model that recapitulates the full progression of autoimmunity-driven tumorigenesis, from gastritis to invasive gastric adenocarcinoma, analogous to the Correa cascade^23^. Although the CTLA4KD model mimics gastric cancer risk in patients with CTLA4 insufficiency, the inflammatory mechanisms underlying tumorigenesis in this model are broadly applicable to gastric cancer development in general^22^. This model offers a unique opportunity to capture the persistence and evolution of molecular and cellular alterations across the entire progression from premalignancy through cancer.

Recent single-cell analyses of gastric metaplasia have begun to define the transcriptional landscape of premalignant SPEM and to identify early markers of the metaplastic transition^24, 25^. However, whether these early changes persist into cancer remains largely unaddressed. In this study, we sought to identify single-cell profiles that arise at premalignancy and persist or evolve through malignancy. Using single-cell transcriptomics of the CTLA4KD mouse gastric mucosa, we identified a metaplastic/tumorigenic cell population that emerges in premalignancy and persists through invasive adenocarcinoma with progressive changes in fibronectin signaling. Experimental validation revealed upregulation of Embigin (EMB), a fibronectin receptor^26^, at both mRNA and protein levels in the gastric mucosa, beginning at the premalignant stage and remaining elevated after cancer development. EMB^+^ cells co-expressed the cancer stem cell marker CD133, linking EMB to a stem-like epithelial phenotype. EMB was also elevated in human gastric adenocarcinoma. These findings present EMB as a candidate marker of gastric premalignant metaplasia and early-stage gastric tumorigenesis.

## Materials and Methods

### Mice

We used male and female mice, as no sex differences were observed. CTLA4KD mice were previously described^22^. Mice used in the study were on a BALB/c x C57BL/6 F1 (CB6F1) genetic background. Gastric mucosa from 2 mice (1 male and 1 female) was pooled for each of 4 samples: Wild-type (WT) and CTLA4KD. Young mice were 5 months old, and aged mice were ∼24-28 months old. The old control samples included 1 CTLA4KD-negative female and 1 PL4 vector transgenic control expressing no CTLA4shRNA. The Institutional Animal Care and Use Committee at the University of Miami approved the studies.

### Cell isolation for single-cell RNA sequencing

Stomachs were cut at the gastroesophageal junction and pyloric sphincter. The forestomach was removed, and tissues were rinsed with PBS. Gastric mucosa was collected by cutting parallel to the mucosal surface, minced into 1-2 mm pieces, and suspended in 5 mL of RPMI with 2% FBS. Tubes were centrifuged at 200 x g for 30 seconds. Media was replaced with 10 mL EDTA dissociation solution (10% FBS, 9% HBSS, 5 mM EDTA, 150 μL 10X HEPES in PBS) and incubated for 15 min on a shaker at room temperature. Tissues settled by gravity (2 min), and the supernatant (gastric crypts and leukocytes) was saved in 5mL of wash buffer (PBS with 0.5% BSA). Remaining tissue was resuspended in 2mL of prewarmed digestion buffer containing 0.8 mg/mL of dispase (Sigma-Aldrich), 0.2 mg/ml of collagenase P (Sigma-Aldrich), and 0.1 mg/mL of DNase I from Worthington Biochemical Corporation (Lakewood, NJ) in RPMI with 2% FBS and incubated at 37° C for 5 minutes. Tissue was mechanically dissociated by pipetting (∼10 times). Supernatant was collected and pooled with the crypt/leukocyte fraction. This digestion process was repeated until all tissue was digested. Samples were centrifuged at 200 x g for 10 minutes, washed in PBS, resuspended in prewarmed trypsin, and incubated at 37° C, 5% CO_2_ for 3 minutes. Samples were passed through an 18G needle, filtered through a 40 μm filter into 10 mL of wash buffer, and centrifuged as above. Samples were resuspended in 5 mL wash buffer, filtered, and centrifuged for 5 minutes at 450 x g. Dead cell removal was conducted by resuspending the pellet in 500 μL of Annexin V binding buffer (BioLegend) with biotinylated Annexin V at a 1:20 dilution. Cells were incubated at 4° C for 10 minutes, washed with 5 mL wash buffer and centrifuged at 450 x g for 5 minutes. Pellet was resuspended in 500 μL wash buffer and 10 μL streptavidin magnetic beads (Stemcell Technologies) and incubated at 4° C for 10 minutes. Sample was resuspended in 5mL wash buffer and centrifuged as above. Pellet was resuspended in 2.5 mL wash buffer, transferred to a 5mL round-bottom polystyrene tube, and placed in an EasySep Magnet (Stemcell Technologies) for 10 minutes. Without removing the tube from the magnet, the contents were poured into a new tube and submitted for sequencing.

### scRNA-seq processing, clustering, and cell-type annotation

Sequencing was conducted at the University of Miami Onco-genomics Shared Resource. We processed raw sequencing data using Cell Ranger 4.0.0 with the mouse mm10 reference genome (mm10-2020-A). Low-quality cells were excluded based on high mitochondrial content, low gene/UMI counts, or extremely high feature counts. After quality control, we merged the samples and analyzed them using Seurat’s SCTransform workflow^27^. Gene expression values were normalized using the SCTransform workflow, and samples were integrated for joint downstream analysis. Principal component analysis was performed for dimensionality reduction, followed by graph-based clustering and UMAP visualization. Cell-type annotation was performed using both SingleR^28^ and AUCell^29^. SingleR was applied with mouse reference datasets, including the Immunological Genome Project, for reference-based predictions. AUCell was used to calculate cell-type signature activity scores based on curated marker-gene sets. Final cell-type identities were assigned by integrating Seurat cluster structure, SingleR predictions, AUCell scores, and canonical marker-gene expression patterns.

### Pseudotime trajectory analysis using Monocle3

Monocle3^30^ cell-data-set objects were constructed from the Seurat-annotated expression matrix and metadata, and the principal graph was learned to order cells along pseudotime. Two separate trajectory analyses were performed, rooting on chief or mucous neck cells, respectively. Marker-gene-based trajectory analysis was performed using curated genes representing gastric epithelial differentiation, proliferation, immune, endothelial, stromal, and metaplastic/tumorigenic states as mentioned in a previous study^25^.

### RNA-velocity and cell-fate analysis using scVelo and CellRank 2

RNA-velocity^31^ and cell-fate analyses were performed independently for WT Young, CTLA4KD Young, WT Old, and CTLA4KD Old samples and were restricted to Chief, IF+ Chief, and Mucous Neck cells. Velocyto-derived spliced and unspliced transcript counts were analyzed using the scVelo dynamical model to estimate RNA velocity, root and endpoint scores, velocity pseudotime, and latent time. The inferred velocity field was visualized on the Seurat-derived UMAP embedding. The resulting velocity-informed transition matrix was analyzed using GPCCA estimator in CellRank 2^32^, configured to identify three macrostates and calculate cell-fate probabilities. Candidate initial and terminal macrostates were inferred automatically without preassigning a biological root, endpoint, or cell type.

### Cell-cell communication analysis using CellChat

Cell–cell communication was analyzed using CellChat ^33^, with the normalized expression matrix and final cell-type annotations as input. Separate CellChat objects were generated for each experimental group using the mouse ligand–receptor database CellChatDB.mouse. Overexpressed genes and ligand–receptor interactions were identified and projected onto the mouse protein–protein interaction network, and communication probabilities were calculated using a truncated-mean approach with population size considered; low-confidence interactions were filtered before analysis. To characterize FN1 signaling by the metaplastic/tumorigenic cluster, per-interaction communication probabilities with this cluster as sender and receiver were extracted from the FN1 pathway-level tables, retaining only interactions passing CellChat’s permutation test (p ≤ 0.05); the relative contribution of individual FN1 ligand–receptor pairs was determined from the ligand–receptor interaction tables.

### Stomach cell isolation for flow cytometric analysis

Mouse gastric mucosa was collected after euthanasia, cut into pieces <5mm in size and incubated in RPMI media (22400-089, Gibco) with FBS (A5256701, Gibco) and Pen/strep (15140-122, Gibco), containing 1mg/ml collagenase IV (C5138, Sigma-Aldrich) and 10 U/ml DNase I (LS002007, Worthington) for 20 minutes at 37° C on a shaker. Following incubation, the supernatant was collected, replaced, and then incubated for 20 minutes at 37° C. This was repeated twice for a total of three 20-minute incubations. Following the incubations, all collected supernatant and undigested tissues were further mechanically dissociated using an 18G needle, filtered with a 70μm filter, and washed with PBS (14190-144, Gibco).

### Full spectrum flow cytometry

A 40-color spectral flow cytometry panel was used to analyze cells, as previously described^34^. Cells were blocked against nonspecific antibody binding using normal mouse serum (SP-002-VX5, ImmunoReagents), anti-CD16/32 antibody (Clone 2.4G2), BD Horizon Brilliant Stain Buffer Plus (566385, BD) and CellBlox blocking buffer (C001T02F01, Thermo Fisher). Cells were stained with antibodies for cell surface markers, followed by fixation and permeabilization for intracellular staining using Transcription Factor Buffer Set (562574, BD). Dead cells were stained and excluded using Zombie NIR (77184, BioLegend) staining. Doublets were gated out from analysis. Manual gating was done using FCS Express software.

### viSNE and SPADE clustering analysis

Spectral flow cytometry data underwent viSNE dimensionality-reducing analysis on the Cytobank platform using FCS files pre-gated on live cells (Zombie NIR^-^) and nonhematopoietic lineage (CD45^-^) cells using FCS Express software. viSNE analysis transformed FCS data into 2 dimensions using the Barnes-Hut implementation of the t-distributed stochastic neighbor embedding (t-SNE) algorithm. Analysis was performed using proportional sampling, with 7,500 iterations, a perplexity of 30, and a θ of 0.5, following Cytobank user guides and similar procedures reported in a previous study^35^. The following markers were used as clustering channels: CD133, CD31, CD38, DMBT1, E-Cadherin, EMB, EpCAM, MUC2, PDPN.

### Cryopreservation and Immunofluorescence staining

All tissues were preserved in OCT compound, sectioned at 6 µm thickness, and mounted on poly-L-lysine-treated slides. Sections were fixed in acetone, rehydrated with PBS, blocked for nonspecific antibody binding, and stained for 1 hour, followed by three 5-minute washes. Murine tissue was stained with DAPI, GS-II lectin Alexa Fluor 488 (L21415, Thermo Fisher), and anti-EMB PE (12-5839-82, Thermo Fisher) antibody. Images were acquired on a Keyence BZ-X700 microscope and analyzed using QuPath software.

Human gastric adenocarcinoma and adjacent normal tissue biopsies from 5 patients were stained with anti-EMB antibody (13946-1-AP, Proteintech), anti-pan cytokeratin Alexa Fluor 594 (914206, BioLegend), donkey anti-rabbit Alexa Fluor 647 (A31573, Thermo Fisher), and DAPI. Images were taken using Echo Revolve fluorescence microscope and analyzed in Fiji (ImageJ). Background was measured from an unstained control and subtracted from the PanCK and EMB channels; no subtraction was applied to DAPI. All channels were sharpened with an unsharp mask (radius 2, weight 0.5), and identical brightness/contrast settings were applied to all images.

### QuPath immunofluorescence quantification

Murine immunofluorescence images were analyzed in QuPath v0.7 at x10 magnification. Regions of Interest (ROI) were manually annotated, and cell detection was performed using the DAPI channel as reference (pixel size 0.5 µm, sigma 1.5 µm, nucleus area 10-400 µm^2^, intensity threshold 10, cell expansion 5.0 µm. Cells were classified by mean channel intensity thresholds applied independently to each channel. Three non-overlapping images per tissue were averaged for analysis.

### TCGA data analysis

RNA sequencing data for the stomach adenocarcinoma cohort (TCGA-STAD) were downloaded from the Genomic Data Commons (GDC) Data Portal (data release v46.0), filtered by project and transcriptome profiling data category. Expression values (TPM) for *EMB* (ENSG00000170571) and *FN1* (ENSG00000115414) were taken from GDC STAR-Counts files aligned to GRCh38. Tumor (n=412) and adjacent normal (n=36), all from TCGA, were analyzed independently using unpaired Mann-Whitney test.

### Statistics

We conducted Student’s t-tests or Mann–Whitney tests (on count data) for single comparisons, and ANOVA with Šidák post-hoc correction for multiple group analyses. When analyzing parametric single comparisons with unequal variances, t-tests with Welch’s correction were used. When analyzing nonparametric multiple groups, Kruskal–Wallis tests were performed with false discovery rate adjustment.

## Results

### scRNA-seq analysis of CTLA4KD gastric mucosa reveals a metaplastic/tumorigenic cluster present in premalignant lesions and persisting in invasive adenocarcinoma

To survey the transcriptomic changes that occur throughout gastric tumorigenesis, single-cell suspensions were made from the gastric mucosa of CTLA4KD mice with premalignant lesions (CTLA4KD Young, ∼5 months of age) and invasive adenocarcinoma (CTLA4KD Old, ∼24-28 months of age), along with age-matched controls (WT Young and WT Old). scRNA-seq data were clustered using the Seurat R package, and clusters were manually annotated based on gene expression (Fig. 1A). We identified resident gastric mucosa populations, including mucous neck cells (*Muc6)*, chief cells (*Chia1*, *Pla2g1*b), and parietal cells (*Atp4a* and *Atp4b)*. Neuroendocrine cells were identified by expression of *Chga* and *Chgb*, and G cells by expression of *Gast*. Stromal clusters were defined by *Col3a1* and *Fn1*, and endothelial cells by *Flt1* and *Ptprb*. Additionally, we identified a metaplastic/tumorigenic cluster expressing metaplasia-and cancer-associated genes, including upregulation of *Aqp5*, *Ccn1, Klf6*, *Muc6*, *Prom1,* and *Sox9,* and downregulation of Ckb, Gkn1, Gkn2, and Tff1 (Fig. S1A, S1B). This cluster was a substantial population in premalignant lesions of young CTLA4KD mice but was nearly absent in WT controls, and persisted in invasive adenocarcinoma (Fig. 1B, S1C), consistent with a transformed cell population.

**Figure 1:**
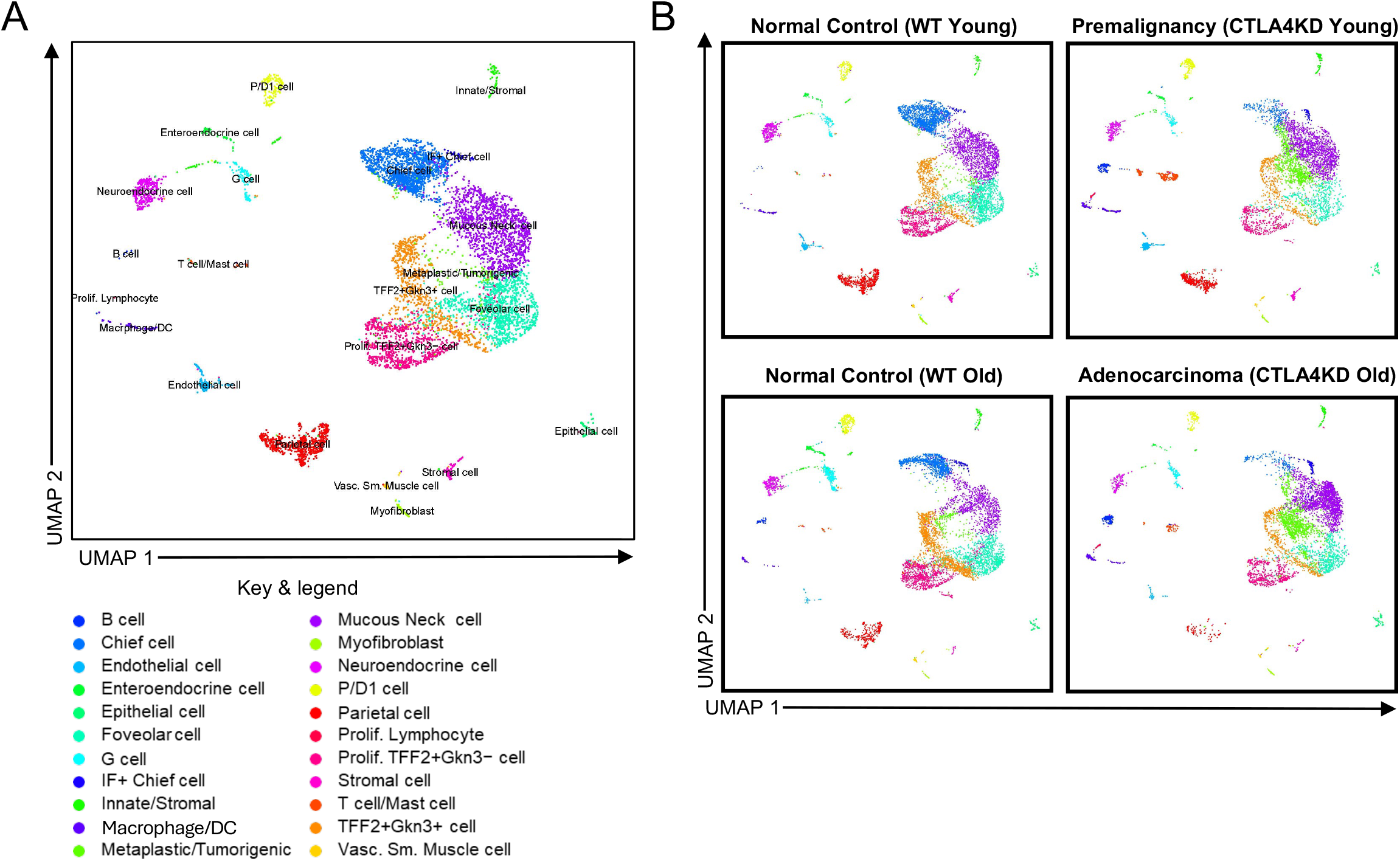
Single-cell RNA-sequencing of gastric mucosa identifies stromal, epithelial, and immune cell populations. Gastric mucosae of wild-type (WT) and CTLA4-knockdown (CTLA4KD) mice at young and old ages were enzymatically dissociated and processed for single-cell RNA-sequencing (scRNA-seq). Cell populations were clustered using the Seurat package in R. (A) UMAP with clusters colored by cell type (see key). (B) UMAPs split by sample: Normal Young Control (WT Young), Premalignancy (CTLA4KD Young), Normal Aged Control (WT Old), and Adenocarcinoma (CTLA4KD Old).

Since our study was performed with single-cell preparation from whole gastric mucosal tissue dissected from the mouse stomach, rather than with isolated gastric glands^24, 25^, we were able to identify major subsets of hematopoietic cells. We identified a macrophage/dendritic cell (DC) cluster (*Itgam*, *Cd84, H2-Aa*), B cells (*Jchain*, *Igkc*, *Igha*, *Ighm*), and a combined T cell/mast cell cluster (*Cd3g*, *Cd3d,* and *Cma1*). The expansion of these hematopoietic clusters in the CTLA4KD mice relative to age-matched controls is expected, as tumorigenesis in this model is driven by autoimmune inflammation (Fig. 1B, S1C).

Together, these data show that our approach of scRNA-seq with whole gastric mucosal tissue captured single-cell profiles from both hematopoietic and nonhematopoietic compartments from premalignancy through cancer. In particular, we identified a cell cluster largely restricted to CTLA4KD mouse mucosa that expresses metaplasia- and cancer-associated genes, which persisted from premalignancy through invasive adenocarcinoma in the CTLA4KD model.

### RNA velocity analysis identifies chief cells rather than mucous neck cells as the origin of metaplastic/tumorigenic cells in the CTLA4KD model

Having identified the gastric cell clusters by scRNA-seq, we next sought to determine which population gave rise to the metaplastic/tumorigenic cluster in CTLA4KD mice. We subjected resident mucosal populations to pseudotime trajectory analysis using the Monocle3 package in R^36^. Prior work has suggested that chief cells or mucous neck cells can serve as the origin of metaplastic and tumorigenic cells, depending on the tumorigenic context^25, 37–40^ and has shown that, following parietal cell loss, chief cells can differentiate toward a mucous neck-like phenotype and then into metaplastic cells^38–40^. Consistent with this, our pseudotime trajectory analysis placed chief and mucous neck cells at the least differentiated end of the trajectory, converging on the metaplastic/tumorigenic cluster (Fig. 2A). This ordering was robust to the choice of root, as rooting the trajectory at either chief cells (Fig. 2B) or mucous neck cells (Fig. 2C) produced the same directional ordering toward the metaplastic/tumorigenic cluster.

**Figure 2:**
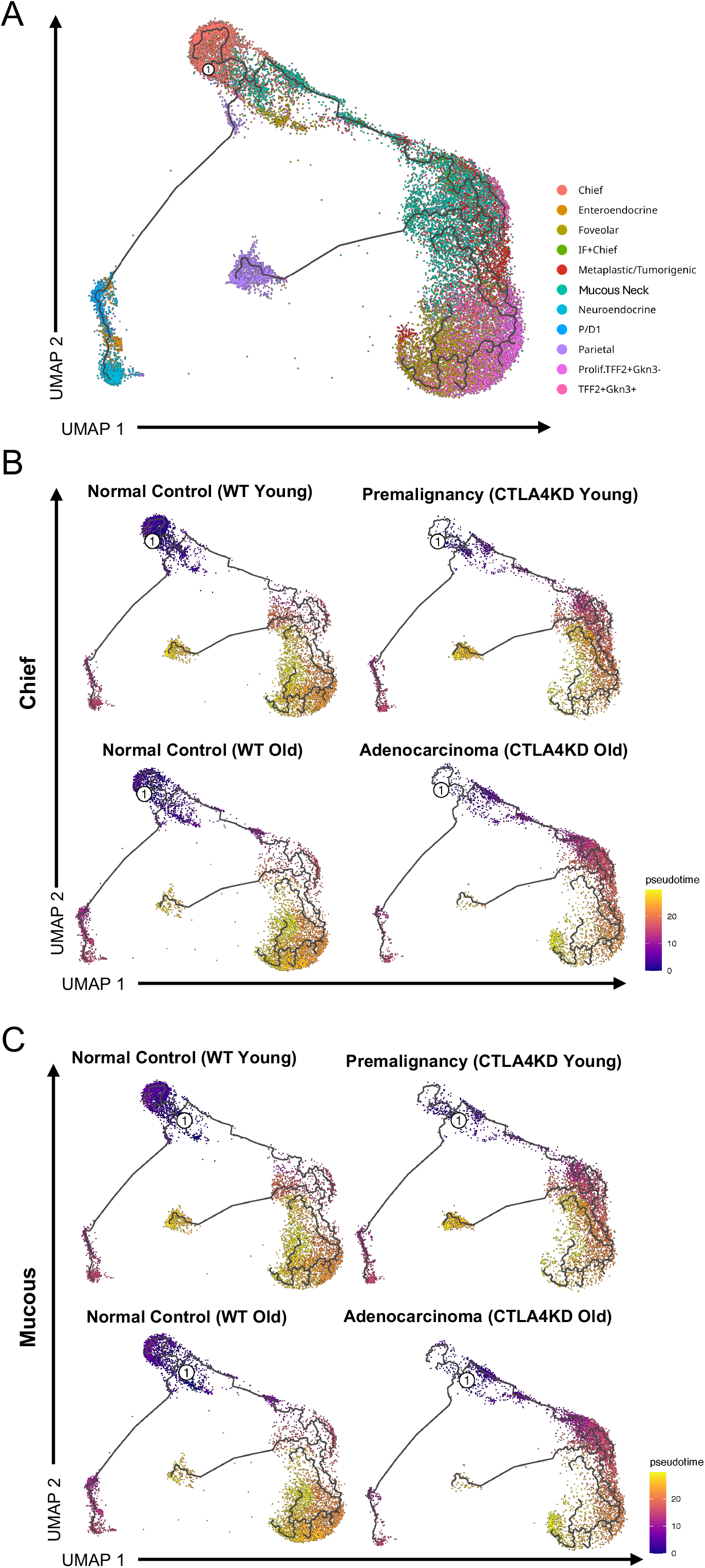
Pseudotime trajectory of non-hematopoietic epithelial lineage cells using Monocle3. Pseudotime trajectory analysis was performed on the scRNA-seq datasets using the Monocle3 package in R. (A) UMAP showing the cell clusters along the pseudotime trajectory (left) with key (right). (B) Pseudotime trajectory rooted at chief cells as the origin population. (C) Pseudotime trajectory rooted at mucous neck cells as the origin population. Purple indicates earlier and yellow indicates later/terminal pseudotime.

To resolve whether chief or mucous neck cell populations give rise to the metaplastic/tumorigenic population without manually selecting a predefined starting point, we applied RNA velocity analysis^31^ (scVelo), followed by CellRank 2^32^, restricting the analysis to chief, IF+ Chief, and mucous neck cells so that directionality among these populations could be assessed independently of the metaplastic cluster. These analyses use the ratio of unspliced to spliced transcripts to estimate the direction in which cells are transitioning and identify velocity pseudotime, latent time, and likely origin and endpoint populations^41^. RNA velocity showed a directional flow from chief cells toward mucous neck cells (Fig. 3A, 3B). Moreover, CellRank 2 identified chief cells as the predicted initial population and mucous neck cells as the terminal population (Fig. 3C). These trajectories were further supported by scVelo outputs, with velocity pseudotime and latent time increasing along the transition from chief to mucous neck cells. Similarly, inferred root cells were enriched in the chief cell cluster, whereas endpoints were enriched within the mucous neck cell population (Fig. S2A-D). Taken together, these analyses suggest a directional transcriptional progression in which chief cells are the most upstream population of the metaplastic/tumorigenic cluster in inflammation-driven gastric tumorigenesis.

**Figure 3:**
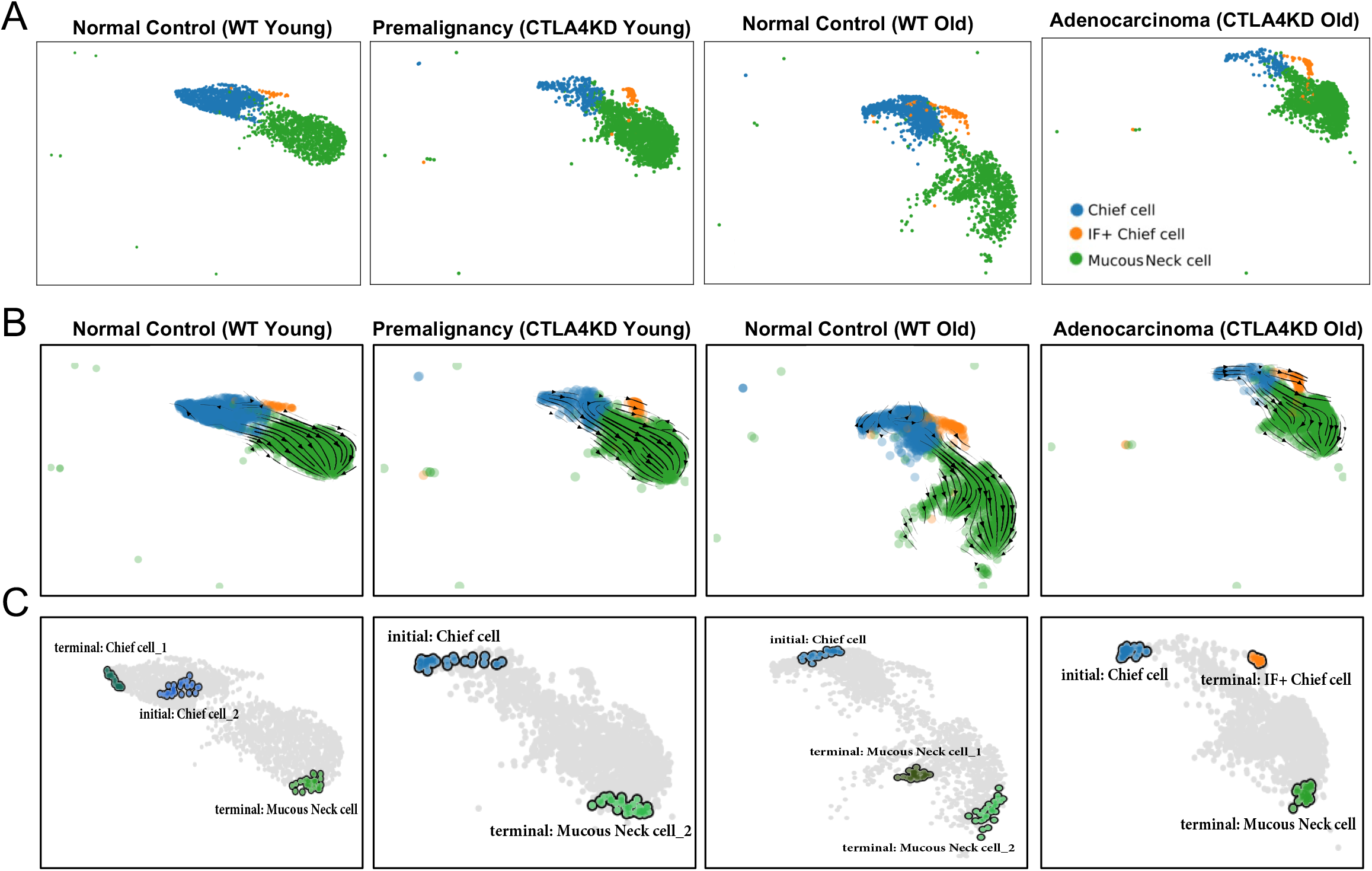
RNA-velocity analysis identifies chief cells as the origin of the metaplastic lineage. RNA velocity analysis was performed on the scRNA-seq datasets using the scVelo package. Analysis was restricted to chief, IF+ chief, and mucous neck cells. (A) RNA velocity-based UMAPs separated by sample. (B) RNA-velocity streamlines projected onto the UMAP; arrows indicate the direction of predicted cell transitions. (C) Automatically inferred initial and terminal states from GPCCA analysis separated by sample.

### Fibronectin (FN1) signaling by the metaplastic/tumorigenic cluster intensifies during tumorigenesis

To identify signaling pathways altered across premalignant and malignant stages, we performed cell-cell communication analysis on our scRNA-seq dataset using CellChat. Among the pathways examined, fibronectin (FN1) signaling was enriched in the metaplastic/tumorigenic cluster and increased progressively across disease stages (Fig. 4A). Expression of *Fn1* was elevated mainly in the stromal cell cluster in both young and old WT controls. However, in CTLA4KD young and old mice, we observed an increase in *Fn1* expression in additional resident gastric cell populations, including foveolar cells, mucous neck cells, proliferating TFF2^+^Gkn3^-^ cells, and the metaplastic/tumorigenic cluster (Fig. S3A).

**Figure 4:**
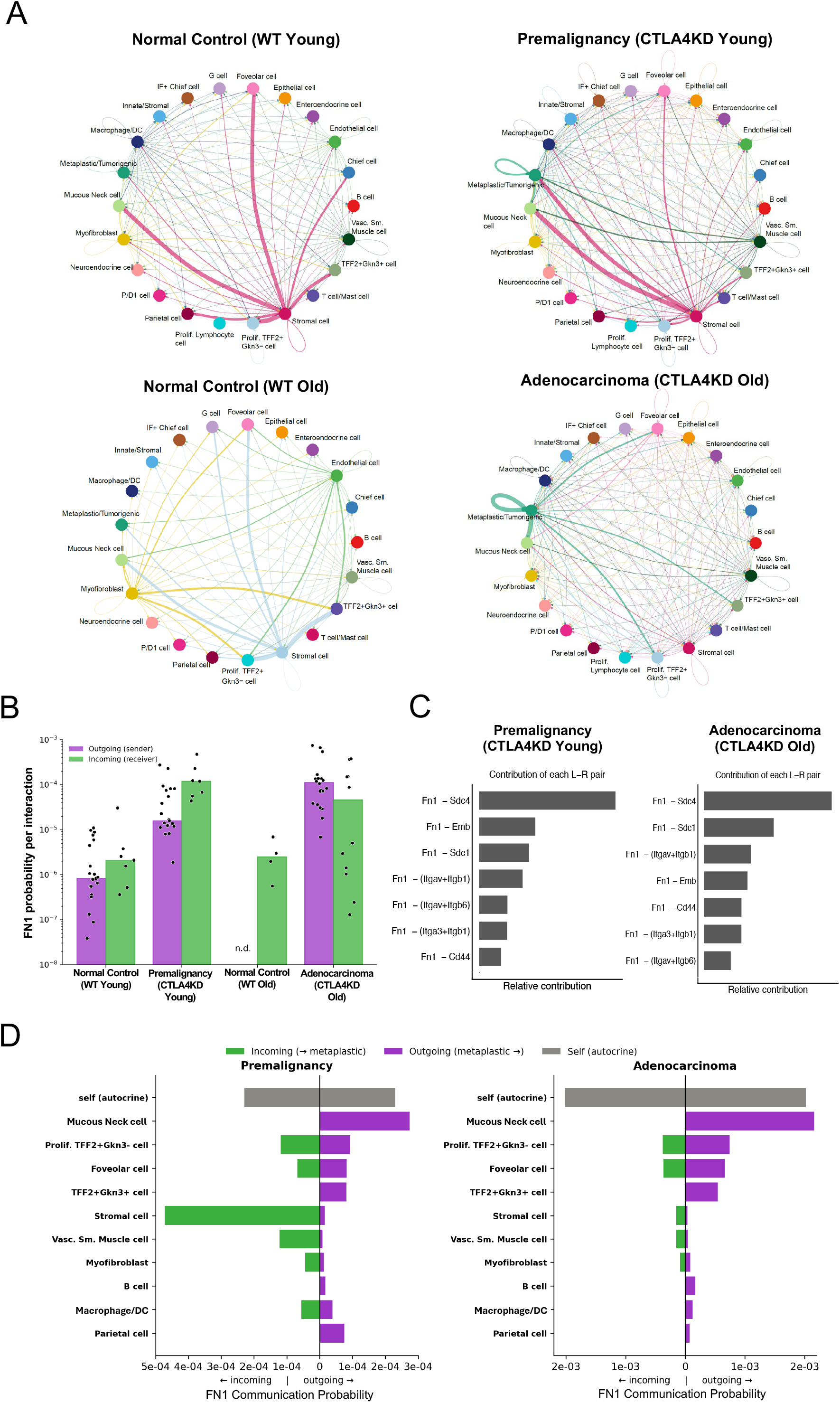
Fibronectin (FN1) signaling by the metaplastic/tumorigenic cluster intensifies during CTLA4KD tumorigenesis. CellChat cell-cell communication analysis of FN1 signaling in the gastric mucosa of WT and CTLA4KD mice at young and old ages. (A) Chord diagrams of predicted FN1 signaling across all cell populations in each sample; edge color denotes the sending population, and edge width denotes the interaction strength. (B) FN1 communication probability per interaction for the metaplastic/tumorigenic cluster as sender (outgoing, purple) and receiver (incoming, green). Each dot represents a significant interaction between the metaplastic/tumorigenic cluster and another cell population; bars denote the median; n.d., not detected. (C) Relative contribution of FN1 ligand-receptor pairs in premalignant (CTLA4KD Young, left) and adenocarcinoma (CTLA4KD Old, right) mucosa (all pairs, Fig. S4A). (D) FN1 signaling partners of the metaplastic/tumorigenic cluster in premalignancy (CTLA4KD Young) and adenocarcinoma (CTLA4KD Old); incoming (green), outgoing (purple), and autocrine (self, gray). Significance by CellChat permutation test, p ≤ 0.05.

Given fibronectin’s role in promoting tumor invasion, epithelial reprogramming, and metastasis in multiple carcinomas^42, 43^, we next examined FN1 signaling at the level of individual cluster interactions. The metaplastic/tumorigenic cluster engaged a similar number of significant FN1 interactions throughout the premalignant and adenocarcinoma stages (21 and 22 as a sender, respectively), yet the median communication probability per interaction increased approximately 7-fold from premalignancy to cancer (Fig. 4B). The escalation in FN1 signaling may reflect an increase in the strength of existing interactions rather than the recruitment of additional partners. Further analysis showed that FN1 signaling in both premalignancy and adenocarcinoma stages appeared to be mediated mainly through syndecans (Sdc4, Sdc1), CD44, and integrin receptors (Itga5-Itgb1, Itgav-Itgb1), as well as Embigin (Emb), a recently identified fibronectin receptor^26^ (Fig. 4C, S4A).

To examine the cellular populations involved in this signaling pathway, we compared the FN1 sources and targets of the metaplastic/tumorigenic cluster across disease stages (Fig. 4D). At the premalignant stage, the metaplastic/tumorigenic cluster received FN1 signaling from stromal cells, vascular smooth muscle cells, and proliferating TFF2^+^Gkn3^-^ cells, as well as autocrine signaling. By the adenocarcinoma stage, incoming signaling to the metaplastic/tumorigenic cluster from the stromal cell population diminished, whereas autocrine self-signaling became the dominant FN1 interaction, exceeding signaling from all other cells (Fig. 4D, S4B). During both premalignant and malignant stages, the metaplastic/tumorigenic cluster also signaled outward predominantly to surrounding metaplastic and mucous neck cells (Fig. 4D). These observations from the computational studies suggest that the metaplastic/tumorigenic population transitions from stroma-dependent fibronectin signaling to a self-sustaining autocrine state as disease progresses.

### Upregulation of the fibronectin receptor Embigin mRNA and protein in gastric premalignancy and adenocarcinoma in the CTLA4KD model

The involvement of Emb in fibronectin signaling in the metaplastic/tumorigenic cluster prompted us to examine the role of this recently described fibronectin receptor further. We found that Emb expression was increased in the metaplastic/tumorigenic cluster in premalignancy (CTLA4KD Young), and adenocarcinoma (CTLA4KD Old) compared with age-matched WT controls (Fig. 5A). Emb can function as a chaperone protein for monocarboxylate transporters^26, 44–46^, so its upregulation could reflect either adhesion or metabolic remodeling. The single-cell profiles of this cluster also exhibited elevated expression of *Prom1* (CD133) compared to WT controls (Fig. 5B). CD133 is a well-established cancer stem cell marker that has been implicated in tumorigenesis, metastasis, and resistance to apoptosis^47–49^.

**Figure 5:**
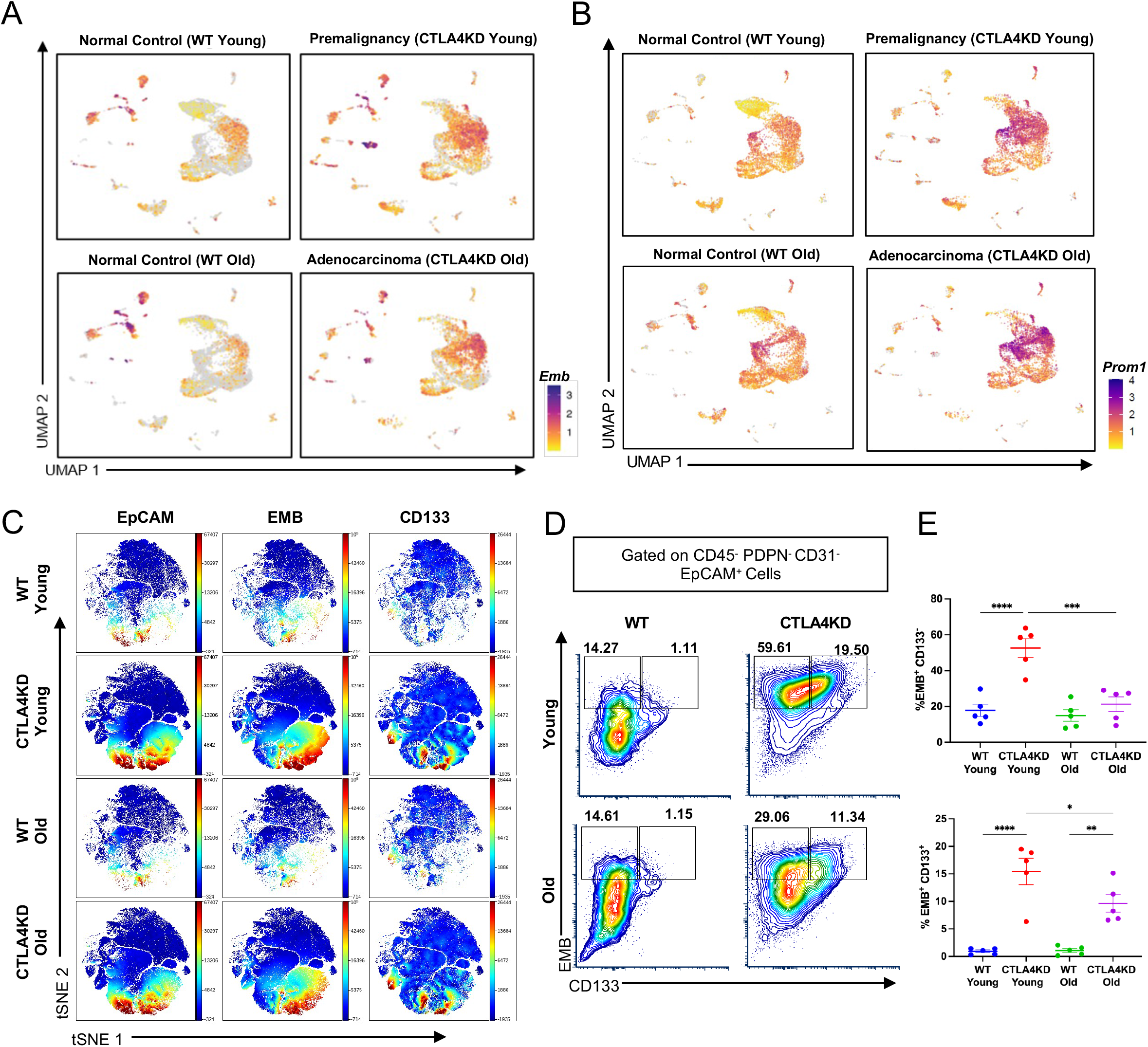
EMB marks a stem-like cell population in the CTLA4KD gastric mucosa. (A) UMAPs showing *Emb* expression. (B) UMAPs showing *Prom1* expression. Protein expression was assessed by spectral flow cytometry of enzymatically dissociated gastric tissues from young and old WT and CTLA4KD mice, followed by viSNE dimensionality-reduction analysis. (C) viSNE plots of non-hematopoietic gastric cells showing EpCAM, EMB, and CD133 expression; blue indicates low expression and red indicates high expression. (D) Representative flow cytometry plots of EMB and CD133 expression with % gated shown. (E) Quantification of the proportion of EMB^+^CD133^-^and EMB^+^CD133^+^ cells (mean ± SEM; n=5; *p ≤ 0.05, **p ≤ 0.01, ***p ≤ 0.001, ****p ≤ 0.0001).

Given that *Prom1* was among the genes defining this cluster (Fig.1), we next evaluated both EMB and CD133 at the protein level. We dissociated gastric mucosa from WT and CTLA4KD mice at young and old stages and analyzed it by 40-color spectral flow cytometry, followed by viSNE dimensionality reduction analysis on the Cytobank platform. Epithelial cells were identified as CD45^-^PDPN^-^CD31^-^EpCAM^+^. viSNE analysis identified a cluster of EpCAM^+^EMB^+^ cells that also expressed CD133 (Fig. 5C). Within the epithelial compartment, EMB^+^CD133^-^ cells were increased in CTLA4KD Young relative to WT Young mice but were comparable to controls in old mice, declining significantly between the premalignant and adenocarcinoma stages. In contrast, EMB^+^CD133^+^ cells were increased in CTLA4KD mice at both ages compared with age-matched WT controls (Fig. 5D, 5E).

Together, these results demonstrate EMB upregulation at the mRNA level within the metaplastic/tumorigenic cluster and at the protein level within the epithelial compartment of CTLA4KD mice, beginning at the premalignant stage and persisting through adenocarcinoma. Notably, the EMB^+^CD133^+^ subset remained elevated through adenocarcinoma, whereas the EMB^+^CD133^-^ population did not, indicating that the CD133-coexpressing population may be the epithelial subset that sustains tumorigenic progression.

### EMB colocalizes with GS-II in the gastric mucosa of CTLA4KD mice

To directly visualize EMB expression in the gastric mucosa, we performed immunofluorescence staining and quantified EMB^+^ cell density across groups using automated cell classification in QuPath. EMB^+^ cell density significantly increased during the premalignant stage (CTLA4KD Young) relative to age-matched WT controls and remained significantly elevated at the adenocarcinoma stage (CTLA4KD Old) relative to age-matched WT controls (Fig. 6A, 6B). These findings confirm at the tissue level that increase in EMB is detectable as early as the premalignancy stage.

**Figure 6:**
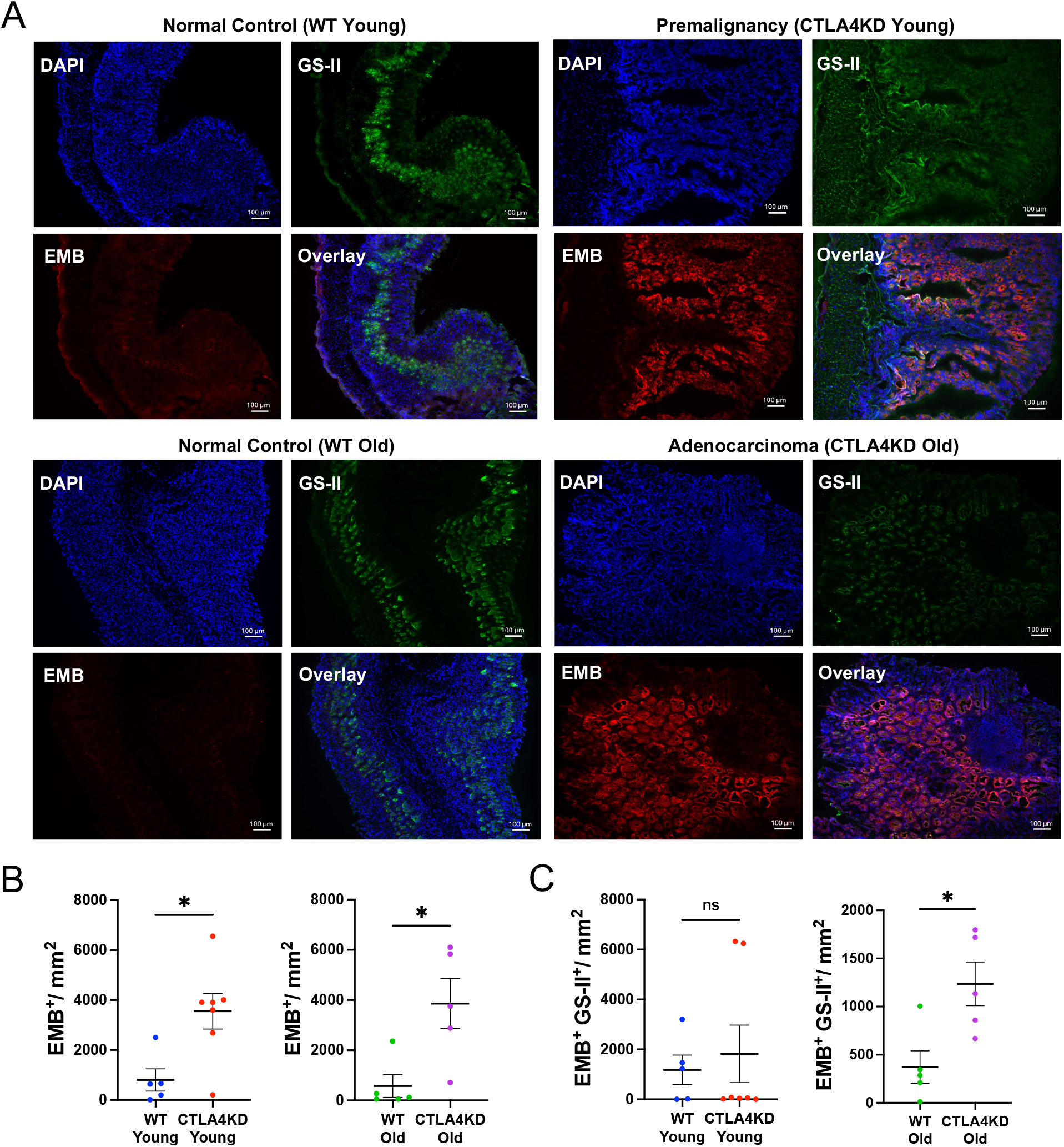
EMB protein expression is upregulated in CTLA4KD gastric mucosa. (A) Representative immunofluorescence images of normal young control (WT Young), premalignancy (CTLA4KD Young), normal aged control (WT Old), and adenocarcinoma (CTLA4KD Old) gastric mucosae stained for DAPI, GS-II, and EMB. (B) Quantification of EMB^+^ cell density (cells/mm^2^ of annotated tissue ROI) in WT Young versus CTLA4KD Young, and in WT Old versus CTLA4KD Old. (C) Quantification of EMB^+^GS-II^+^ cell density (cells/mm^2^ of annotated tissue ROI) in WT Young versus CTLA4KD Young, and in WT Old versus CTLA4KD Old (mean ± SEM; n=5; *p ≤ 0.05). Scale bars, 100 µm.

We next examined colocalization of EMB with *Griffonia simplicifolia* lectin II (GS-II), a lectin that marks mucous neck cells and metaplastic gastric epithelium. The density of EMB^+^GS-II^+^ cells was significantly increased in the CTLA4KD Old mice relative to age-matched WT controls but did not differ between CTLA4KD Young and WT Young mice (*P=0.92*; Fig. 6C). In other words, although EMB upregulation was present from the premalignant stage, its colocalization with the GS-II^+^ metaplastic compartment was detected only at the adenocarcinoma stage, suggesting a dynamic progression of EMB^+^ population as disease advances.

### EMB mRNA and protein are detected in human gastric adenocarcinoma

To assess the translational relevance of EMB to human gastric cancer, we analyzed EMB expression in stomach adenocarcinoma and healthy gastric tissue using transcriptomic data from The Cancer Genome Atlas (TCGA). *EMB* mRNA expression was significantly elevated in tumor tissues relative to healthy gastric mucosa (Fig. 7A), consistent with the observation of its upregulation in the CTLA4KD model. *FN1* mRNA expression was similarly increased in tumors relative to healthy gastric mucosa (Fig. 7B).

**Figure 7:**
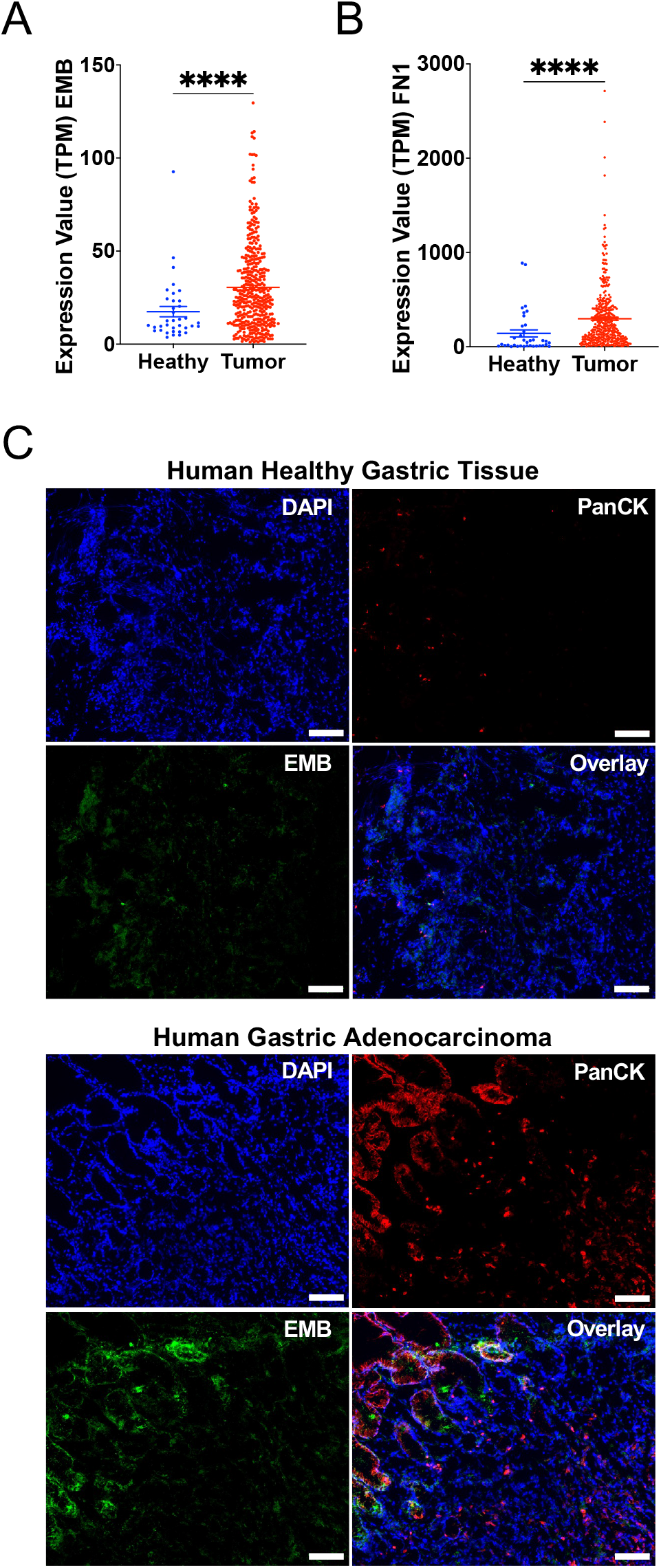
EMB is upregulated in human gastric adenocarcinoma compared with normal gastric tissue. (A) Expression of *EMB* (TPM) and (B) *FN1* (TPM) in primary tumor (n=412) and adjacent healthy (n=36) tissue from TCGA-STAD cohort. (C) Representative immunofluorescence images of matched human healthy gastric tissue (top) and gastric adenocarcinoma (bottom) stained for DAPI, PanCK, and EMB (mean ± SEM; ****p ≤ 0.0001). Scale bar: 100 µm.

To corroborate these findings, we obtained matched gastric adenocarcinoma and adjacent healthy tissue biopsies from male and female donors aged 31 to 66 years. We performed immunofluorescence staining for EMB protein, the epithelial marker pan-cytokeratin (PanCK), and DAPI. Across the patient samples, gastric adenocarcinoma tissues showed stronger staining of EMB than healthy gastric tissue, with EMB signal colocalizing with PanCK^+^ epithelial regions in tumor samples (Fig. 7C, S5). Although limited to a small number of donors (n=5), the consistent pattern across samples suggests that EMB upregulation is shared between the CTLA4KD model and human gastric adenocarcinoma, supporting its potential as a translationally relevant marker of gastric tumorigenesis.

## Discussion

Early detection of gastric cancer remains a major clinical challenge, given that symptoms typically begin after disease has progressed to advanced stages. The molecular changes that distinguish premalignant metaplasia from normal gastric mucosa, and that could enable earlier detection, remain poorly defined. Here, we delineated single-cell transcriptomic profiles from premalignancy through invasive adenocarcinoma and identified candidate markers that can potentially be used in combination to reveal a subset of cancer-progenitor cells and their pathways to invasive malignancy. In-depth computational analysis and experimental validation identified EMB, a recently identified fibronectin receptor, as a marker upregulated in the premalignant gastric mucosa that persists through malignancy.

To capture the transition from premalignancy to cancer, we profiled the gastric mucosa of CTLA4KD mice, which develop autoimmune inflammation that drives the entire tumorigenic cascade from the early stage of premalignancy (SPEM) to invasive adenocarcinoma. Single-cell transcriptomics revealed major immune cell subsets in the inflammatory setting, including T cells, B cells, and innate immune cells, along with resident gastric mucosa cells including parietal, chief, mucous neck, and stromal cells. Importantly, this study identified a metaplastic/tumorigenic cluster that persists and evolves in the setting of autoimmune inflammatory tumorigenesis in the stomach.

The majority of gastric cancer cases are originally triggered by *H pylori* through a well-described process ^5, 50, 51^. Autoimmune-driven gastric tumorigenesis is thought to follow a similar histopathological sequence and inflammatory pathways ^8, 22^. Indeed, one study found that H. pylori and autoimmune-driven metaplasia are difficult to distinguish at the cellular level and share metaplastic lineages^52^. Chief, progenitor, and mucous cells have each been implicated as origins of metaplasia^25, 37–40^. To determine which population is most likely to give rise to the metaplastic/ tumorigenic cluster in our CTLA4KD model, we performed standard pseudotime trajectory analysis, in combination with more recently developed approaches, RNA velocity analysis (scVelo) and CellRank 2. The findings from our study support chief cells as the likely cellular origin of the metaplastic/tumorigenic population in our model of autoimmune tumorigenesis. Lineage tracing experiments in the CTLA4KD model will be required to identify the specific cell of origin and to establish its contribution to cellular transformation.

Recent advances in bioinformatic tools also enabled us to analyze the single-cell transcriptomic profile beyond the typical approaches based on differential gene expression, for example, using CellChat ^33^ to infer cell-cell communication. The analysis revealed that fibronectin (FN1) signaling was enriched in the metaplastic/tumorigenic cluster and intensified from premalignancy to adenocarcinoma. Fibronectin is an extracellular matrix glycoprotein produced largely by stromal and mesenchymal cells that mediates cell adhesion, migration, and matrix organization^43^, and its accumulation within the tumor microenvironment has been linked to stemness across multiple carcinomas^53, 54^. Notably, in our model, the source of this signaling shifted during tumorigenesis. At the premalignant stage, the metaplastic/tumorigenic cluster received fibronectin predominantly from stromal populations, whereas at the adenocarcinoma stage, autocrine signaling became the dominant source. This shift from stroma-dependent to self-sustaining fibronectin signaling suggests that these cells progressively establish their own fibronectin-rich environment as disease advances, reducing their dependence on the surrounding stroma and potentially contributing to a malignant and metastatic state.

Consistent with their tumorigenic potential, a subset of metaplastic/tumorigenic population expressed both EMB and CD133, a cancer stem cell marker^48^. The co-expression of EMB and CD133 further supports a link between EMB and a stem-like phenotype and offers insights into its potential function in gastric tumorigenesis. Taken together, these observations suggest a model in which the metaplastic/tumorigenic population exploits a fibronectin-rich environment through EMB. Aided by its fibronectin-binding activity, EMB may mediate the interaction of these cells with the extracellular matrix and help maintain their undifferentiated, stem-like state, consistent with its role in sustaining progenitor identity in other tissues^26^ and with the CD133 co-expression we observed. Through its chaperone function for MCT1/2, EMB may also support the altered metabolism seen in proliferating tumor cells^46^.

Nevertheless, the role of EMB in cancer appears to be cancer-type dependent. In prostate cancer, EMB expression was higher relative to normal tissues and induced NF-κB and MMP9 activity, promoting tumor growth and metastasis^55^. In breast cancer, by contrast, EMB was suggested to be tumor-suppressive, as its depletion increases cell proliferation and migration^56^. In this study, Emb upregulation across premalignant and malignant lesions in gastric mucosa leads us to hypothesize that EMB expression may promote gastric cancer development. Emb functions as a chaperone for monocarboxylate transporters 1 and 2 (MCT1/2)^46^, which transport lactate, pyruvate, and other monocarboxylates across the plasma membrane^44, 45^. In sebaceous glands, EMB acts as a fibronectin receptor whose loss allows progenitor cell differentiation^26^, with Emb deficiency causing developmental delays and dysfunction in the mouse lung and kidney^57, 58^, implicating it in the maintenance of progenitor states. Further studies are needed to understand whether and how EMB influences gastric tumorigenesis, or whether it is merely a marker of tumor progression reflected by its upregulation in both mRNA and protein in premalignancy through cancer.

The rising incidence of early-onset gastric cancer shows the need for new markers for early detection of malignant transformation even at the premalignant stage. The results from this study suggest EMB as a candidate marker. The upregulation of EMB in human gastric adenocarcinoma relative to healthy gastric tissue indicates that its association with gastric tumorigenesis is conserved across species. However, whether EMB is similarly elevated in human premalignant lesions, such as SPEM and IM, remains to be determined. In-depth studies of EMB in premalignant and cancerous microenvironment with large sets of human tissue samples from various gastric cancer subtypes are needed to rigorously assess the potential of EMB as an early detection marker.

## Acknowledgments

We thank the Cytometry and Imaging Shared Resource (CISR) and the Biostatistics & Bioinformatics Shared Resource (BBSR) at the University of Miami Miller School of Medicine for their support. We also thank Dr. Alejandro Villarino and Molly Dalzell for their valuable input and assistance with single-cell RNA sequencing data analysis. The results shown here are in part based on data generated by the TCGA Research Network: https://www.cancer.gov/tcga

---

**Figure S1.**
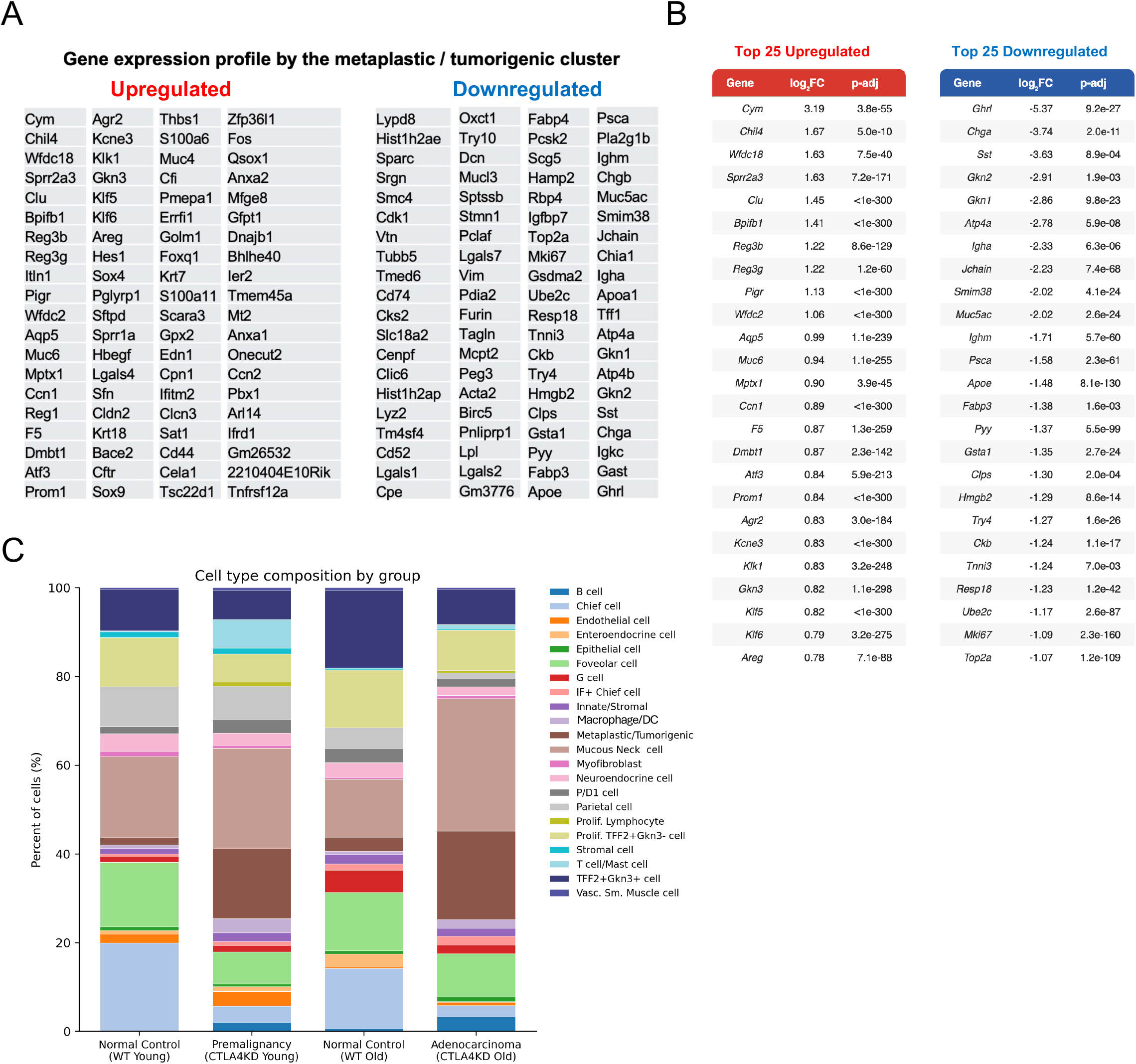
Differentially expressed genes defining the metaplastic/tumorigenic cluster. Differential expression analysis of the metaplastic/tumorigenic cluster in the CTLA4KD gastric mucosa single-cell RNA-seq dataset. (A) Gene expression profile in the metaplastic/tumorigenic cluster. (B) Top 25 upregulated and top 25 downregulated differentially expressed genes in the metaplastic/tumorigenic cluster ranked by average log₂ fold change and adjusted p-value. (C) Stacked bar plot of the relative proportion of each annotated cell population (as a percentage of total cells). Cell proportions for each group are provided in Supplementary File 2.

**Figure S2.**
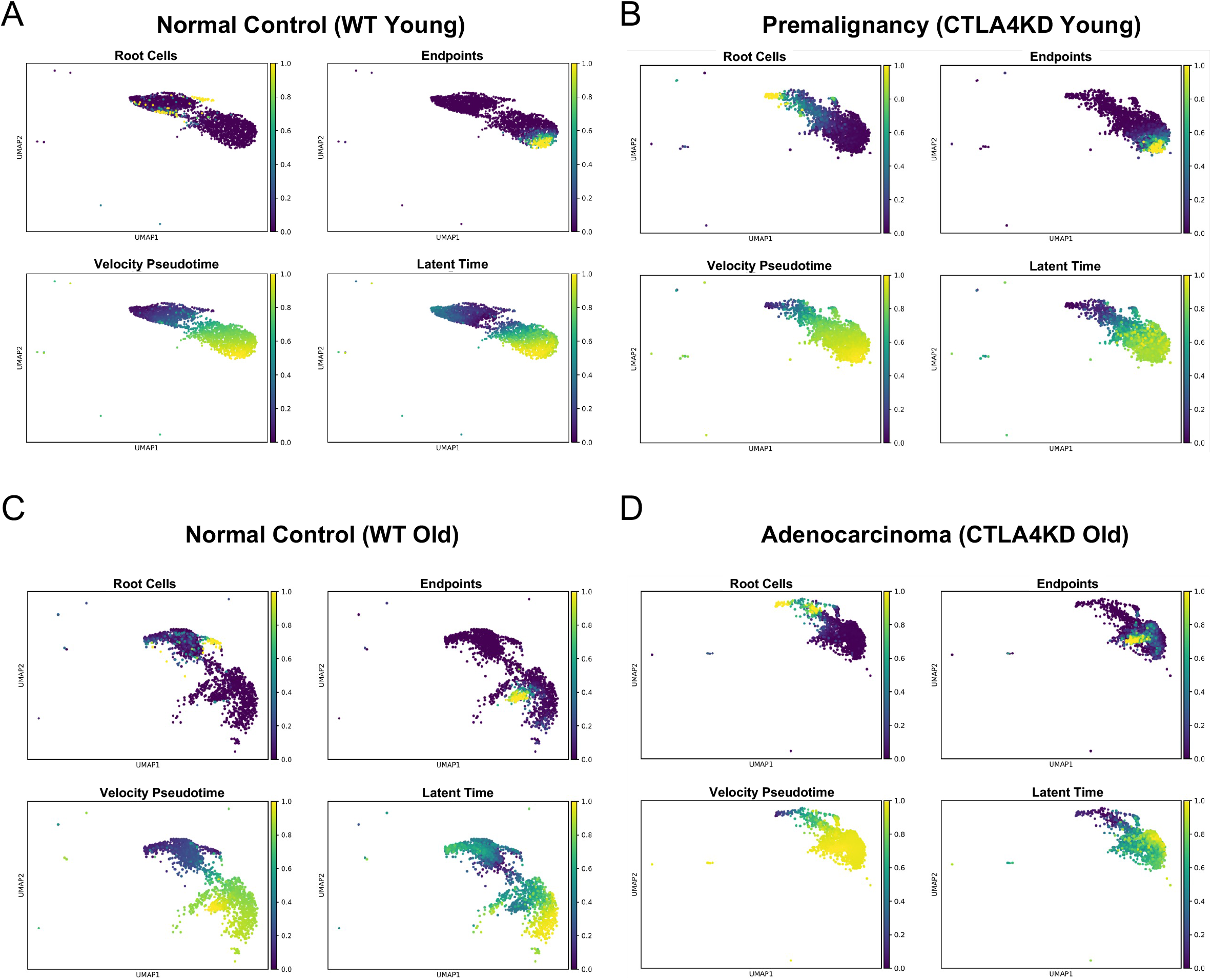
Scores for scVelo analysis of root cells, endpoint cells, velocity pseudotime and latent time. scVelo root cells, endpoints, velocity pseudotime, and latent time UMAPs in (A) Normal Control (WT Young), (B) Premalignancy (CTLA4KD Young), (C) Normal Control (WT Old), and (D) Adenocarcinoma (CTLA4KD Old). Samples were restricted to Chief, IF+ Chief, and Mucous Neck cells.

**Figure S3.**
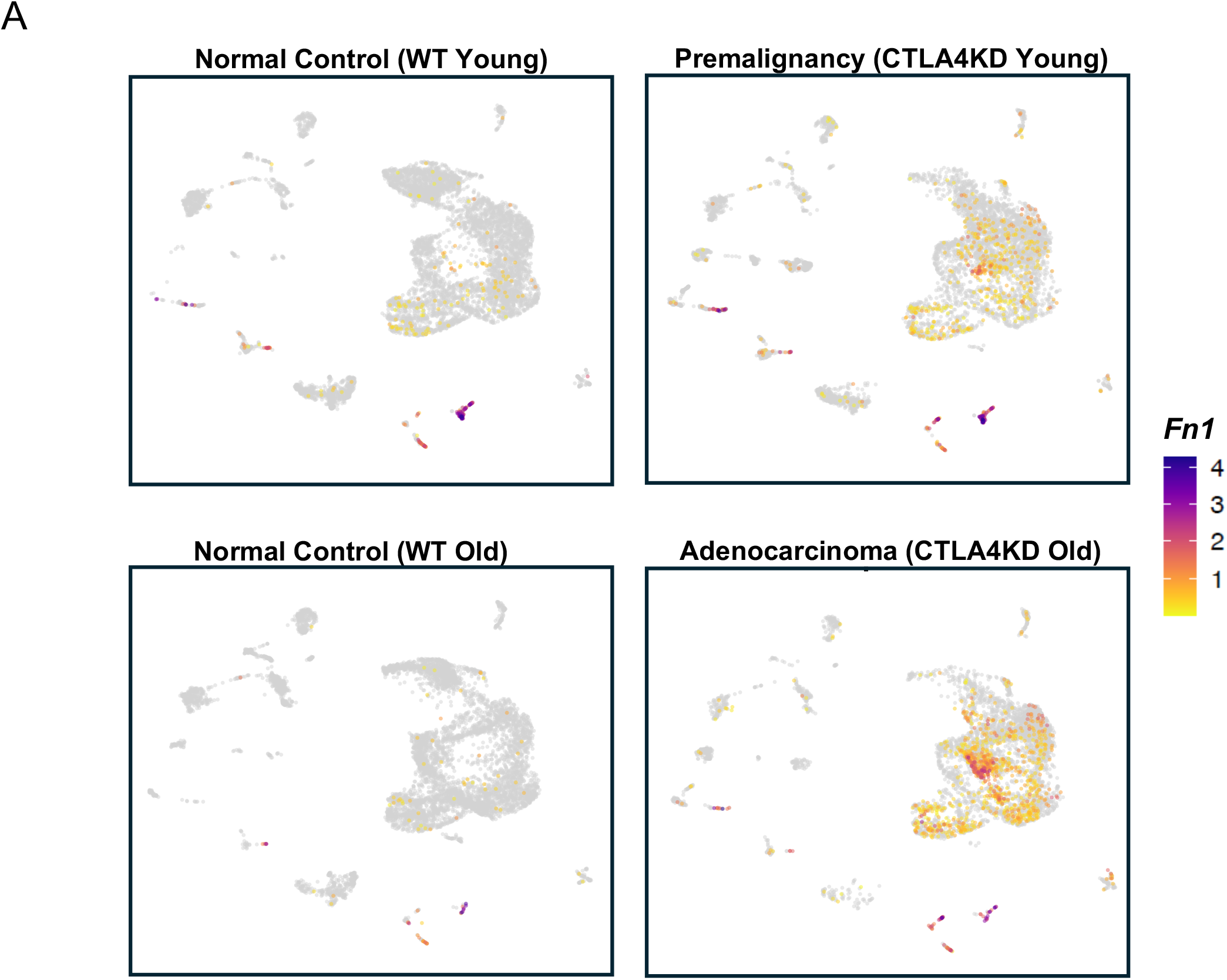
Expression of *Fn1* across groups. (A) UMAPs showing expression of *Fn1,* split by sample: Normal Control (WT Young), Premalignancy (CTLA4KD Young), Normal Control (WT Old), and Adenocarcinoma (CTLA4KD Old).

**Figure S4.**
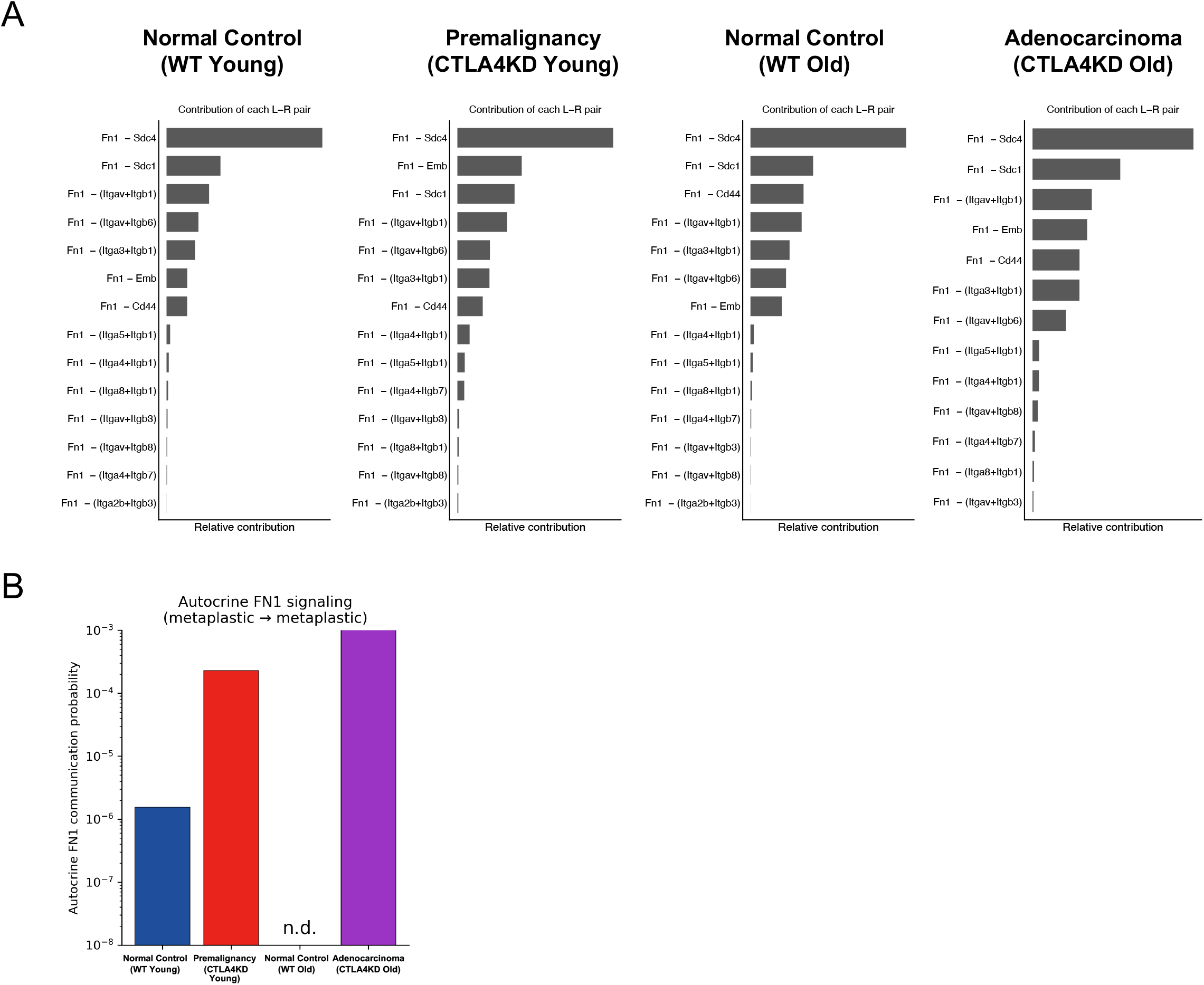
Fibronectin ligand-receptor pair and autocrine FN1 signaling. (A) Relative contribution of individual fibronectin (FN1) ligand–receptor pairs to total FN1 signaling in each group: normal control (WT Young), premalignancy (CTLA4KD Young), normal control (WT Old), and adenocarcinoma (CTLA4KD Old). (B) Autocrine FN1 communication probability (metaplastic/tumorigenic → metaplastic/tumorigenic) across all four groups. n.d. (not detected). Communication probabilities are derived from CellChat; significance was assessed by CellChat permutation test (p ≤ 0.05).

**Figure S5.**
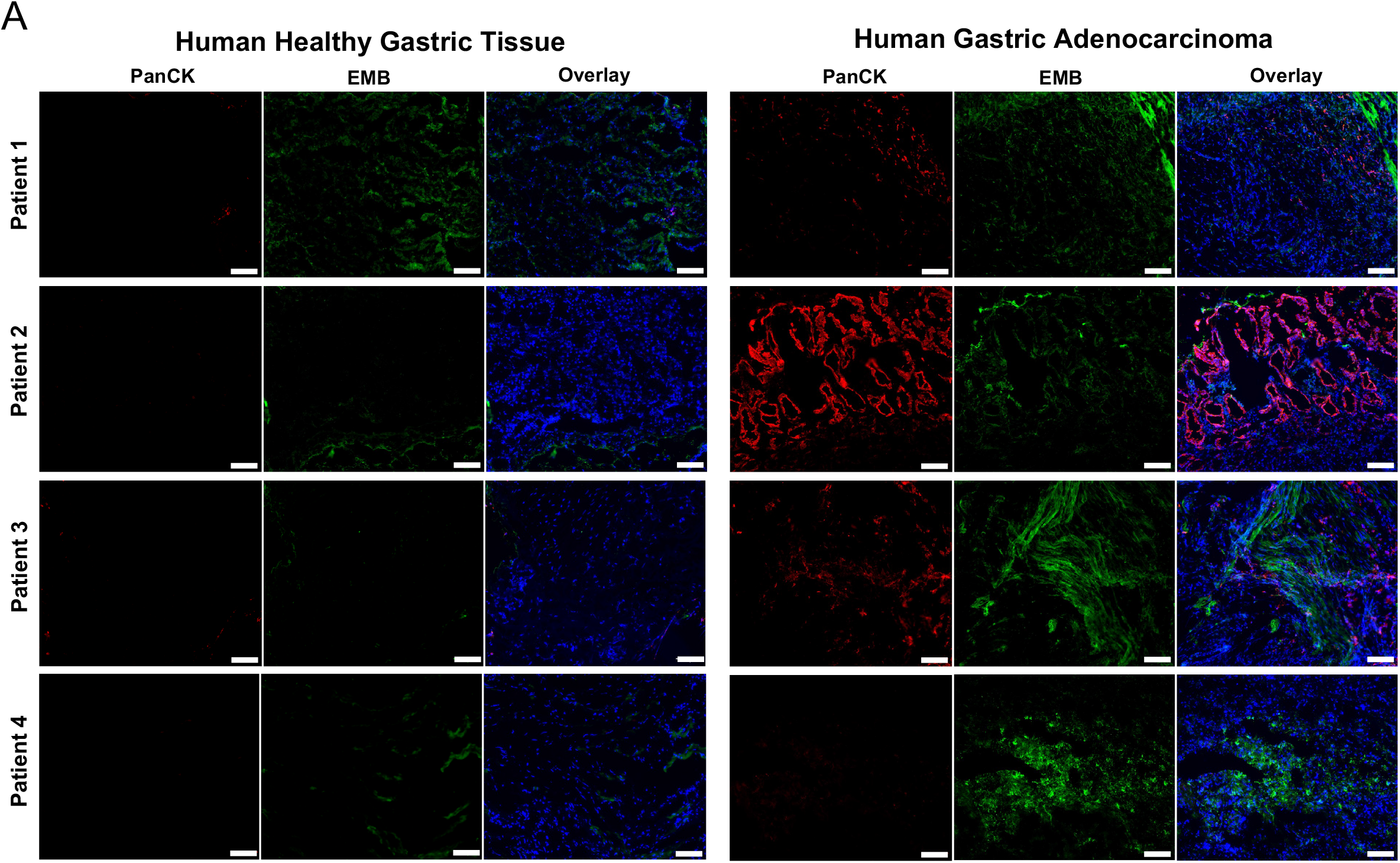
EMB expression in human gastric adenocarcinoma and adjacent healthy tissue. (A) Immunofluorescence images from 4 matched donors; healthy gastric tissue (left) and gastric adenocarcinoma (right) stained for DAPI, PanCK, and EMB. Scale bars: 100 µm; magnification x10.

